# The relationship between auditory brainstem responses, cognitive ability, and speech-in-noise perception among young adults with normal hearing thresholds

**DOI:** 10.1101/2024.12.19.629463

**Authors:** Mishaela DiNino, Jenna Crowell, Ilsa Kloiber, Melissa J. Polonenko

**Affiliations:** Department of Communicative Disorders and Sciences, University at Buffalo, Buffalo, NY 14214, U.S.A; Neuroscience Program, University at Buffalo, Buffalo, NY 14203, U.S.A; Department of Speech-Language-Hearing Sciences, University of Minnesota, Minneapolis, MN 55455, U.S.A

**Keywords:** Young adult, normal hearing, speech-in-noise perception, auditory brainstem response, cognition, evoked responses

## Abstract

The goal of this research was to determine the contributions of auditory neural processing and cognitive abilities to predict performance on a speech-in-noise perception task in young, normal hearing adults. Two experiments were performed, each with separate cohorts of ∼30 young adults with normal hearing who performed a competing talker task which included a high-pass filtered condition that was designed to be more sensitive to auditory nerve functioning than are commonly used speech-in-noise perception tests. Predictors of performance on this competing talker task included ABR waves I and V metrics and cognitive test scores. Experiment one included click ABRs at a moderate level commensurate with the level of the competing talker task, as well as the cognitive digit span working memory test. Experiment two included high-intensity click clinical ABRs and three cognitive tests from the NIH Toolbox V3 that assessed working memory, cognitive flexibility and attention, and inhibitory control: List Sorting Working Memory, Dimensional Change Card Sort, and Flanker Inhibitory Control and Attention tests, respectively. Performance on the high-pass filtered competing talker task varied across participants in both experiments. This variability was predicted by performance on the test of inhibitory control, but not the tests involving working memory or cognitive flexibility, nor by any of the auditory processing metrics from moderate or high-intensity click ABRs. Among two groups of young adults with normal hearing, cognitive factors with very similar demands to the competing talker task seem to play the greatest role in speech-in-noise perception.

## 1. Introduction

Understanding speech presented in background noise depends on both auditory system function and higher-level cognitive abilities. Adequate peripheral and central auditory processing are necessary to accurately perceive target sounds in the presence of competing maskers (Dai et al, 2018). Cognitive factors such as attention, executive function, and working memory also play a substantial role in following conversations in noisy listening environments, allowing a listener to focus on a target talker while ignoring background noise, and to hold sentences in memory before responding. Listeners may even be able to compensate for reduced auditory system function with enhanced cognitive strategies (Gordon-Salant & Cole, 2016). There is much evidence supporting a link between age-related decline in auditory sensory and cognitive ability and older adults’ challenges hearing in noise (Humes & Roberts, 1990; Moore et al., 2014). However, among young adults with normal cochlear function, it is less clear how the subsequent structures in the ascending auditory pathway as well as cognitive factors contribute to speech-in-noise perceptual abilities.

The auditory nerve (AN) preserves and transmits all features of a sound following cochlear processing. A study conducted by Lopez-Poveda and Barrios (2013) in young adults with normal hearing thresholds showed that simulating a reduced AN population response from cochlear deafferentation decreased the faithfulness of auditory signal encoding and increased noise in the auditory representation. This manipulation affected listeners’ sentence recognition in background noise more so than in quiet, demonstrating the importance of optimal auditory nerve processing for speech-in-noise recognition. In quiet, the AN encodes only one signal, but in noise the AN encodes multiple signals that a listener must segregate to achieve accurate perception. Degradation of sound features due to decreased AN function, then, would be expected to interfere with auditory perception in background noise much more so than in quiet.

Research in animal models also provides support for a relationship between early neural processing and hearing in noise. In one such study, rats with normal auditory detection thresholds but reduced AN response amplitude from putative noise-induced cochlear synaptopathy exhibited decreased ability to perceive the sound that served as a warning for a subsequent tactile startle stimulus when background noise was present (Lobarinas et al. 2017). In another study, cochlear deafferentation in mice resulted in poor tone detection in noise but not in quiet. These results stemmed from increased central gain following the lower number of AN fibers able to transmit auditory signals, suggesting that diminished AN function may have cascading effects on higher structures in the auditory neural pathway (Resnik & Polley, 2021). Reduced AN response amplitude, then, may be indicative of both decreased AN function and the potential for other, upstream changes in auditory neural processing that could affect hearing in noise.

However, studies in humans have shown mixed evidence for a relationship between neural processing and speech perception in the presence of competing maskers. Young adults with normal hearing thresholds vary in their auditory brainstem responses (ABRs; Prendergast et al., 2017a; Rowe, 1978), which are electrophysiological measures of AN and subcortical nuclei responses to sound. Yet, lower AN and subcortical function, evidenced by reduced ABR wave amplitudes and/or longer wave latencies, does not always correlate with perceptual deficits in noise (Grose et al., 2017; Prendergast et al. 2017b; Smith et al., 2019). One factor complicating the exploration of this relationship is that the etiology of individual differences in ABRs in the absence of audiometric hearing loss is unknown in humans. ABRs are noninvasive, indirect measures of neural processing and thus the specific cause of reduced ABRs in normal hearing listeners cannot typically be determined. Another factor that may interfere with an examination of the relationship between ABRs and one’s ability to hear in noise is that individuals with reduced auditory neural processing may have compensatory strategies, such as enhanced cognitive ability, that allow them to perform similarly on perceptual tasks as listeners with optimal AN and subcortical function.

Many previous studies have found cognitive task performance to coincide with speech-in-noise perception scores in middle-aged and older adults (Füllgrabe & Rosen, 2016a; Vermeire et al., 2019) and among participant groups with a wide age range and hearing thresholds (Valderrama et al., 2018; Yeend et al., 2017; Yeend et al., 2019). Although older adults tend to exhibit a decline in both cognitive and sensory function, some evidence suggests that those who maintain good cognitive ability are those who perform better on speech identification in background noise (Gordon-Salant & Cole, 2016). Still, previous research is mixed as to whether cognitive domains such as working memory capacity predict performance on speech-in-noise perception tasks among *young* adults with normal hearing thresholds (Füllgrabe & Rosen, 2016b; Tamati et al., 2013). An open question is whether young adults exhibit enough individual differences in cognitive function for it to significantly impact hearing in noise.

In addition, the strength of the relationship between cognitive function and speech-in-noise perception for any age group will depend on the type of speech stimulus used. For example, Heinrich et al. (2015) found a significant relationship between attention and working memory scores and sentence identification in noise, but not digits or phoneme identification in noise. Tests of sentence recognition inherently involve higher-level cognitive abilities, a factor that makes sentence stimuli more akin to real-world listening than tests of digit/phoneme identification. However, using sentence stimuli may obscure any potential relationship between deficits in sensory processing and speech-in-noise perception scores. Listeners who are better able to attend to the whole sentence while ignoring competing sounds and/or hold the sentence in memory until a response is prompted may perform well on the task despite any diminished auditory system function.

A potential solution to the dilemma of whether to use speech perception tasks that have low ecological validity but that assess basic sensory perception vs. implement tests with greater similarity to real-world listening but with higher non-sensory, cognitive demands is to use a sentence recognition test, representative of everyday listening, in which the stimuli have been manipulated to be more sensitive to sensory processing. In this study, we high-pass filtered sentences so that their identification in the presence of competing talkers is more sensitive to AN function than are commonly used speech-in-noise perception tests. The filtering removed the frequencies that produce clear cochlear excitation peaks (the first eight harmonics; Oxenham, 2021), such that listeners needed to primarily rely on pitch coded by AN phase-locking to differentiate between the fundamental frequencies (F0s) of a target talker and a competing talker that are presented from the same spatial location. Reduced AN function, then, should affect a listener’s ability to segregate the talkers and accurately identify the target sentence during this task.

The intensity level of stimuli used to elicit ABRs may also play a role in the relationship between ABR wave amplitudes and latencies and speech-in-noise perception scores. Clinical paradigms use very high intensity clicks to maximize the ability to view each component wave of the response. However, this high intensity is not typical of listening conditions in daily life. ABRs to high presentation levels therefore represent a different range of the nonlinear cochlear response, as well as a larger spread of neural activation, than would be elicited by sounds such as everyday conversational speech. The standard ABR parameters were designed for clinical assessment of auditory system dysfunction and not necessarily for comparison to behavioral task performance. It is yet unknown whether ABRs in response to stimuli presented at the same intensity as real-world listening scenarios, and are thus more representative of the auditory system activation that occurs in those scenarios, would better relate to performance on auditory perceptual tasks.

The goal of this research was to examine the respective and interactive roles of neural processing and cognitive ability on speech-in-noise perception performance among young normal hearing adults. Across two experiments, scores on the high-pass filtered competing talker task were compared to click ABRs in response to stimuli that were presented at levels 1) representative of sounds heard in real-world listening environments, and 2) consistent with those used for clinical diagnosis. Measures of working memory and executive function were also collected to determine cognitive contributions to young adults’ ability to identify a target sentence in the presence of a masking sentence, as well as to examine whether cognitive task performance can offset the consequences of reduced neural function on young adults’ speech-in-noise perception test performance.

## 2. Experiment 1

### 2.1. Materials and Methods

#### 2.1.1 Participants

A total of thirty young adults completed Experiment 1. One participant’s data was excluded due to excessive noise in the ABR recordings. The final sample included 29 participants aged 18-32 years, with an average age of 23.83 years. All participants were verified to have normal hearing thresholds by passing an audiometric screening at 20 dB HL for all octave frequencies between 250 Hz and 8000 Hz. Participants reported no history of neurological disorders or hearing difficulties and reported proficiency in English. All study procedures were approved by the University of Minnesota Institutional Review Board. Participants gave written informed consent at the beginning of the study appointment and were compensated for their time.

#### 2.1.2 Auditory Brainstem Responses

Participants were fitted with an appropriately sized electroencephalography (EEG) cap containing 32 pin-type electrodes in accordance with the 10-20 system. One reference electrode was placed on each earlobe, which results in less recording noise than references placed on the mastoids. Electrode offset voltages were verified to be ≤ 20 mV via continuous monitoring, with adjustment of the EEG cap and/or electrodes as necessary. EEG cap set-up took approximately 15-20 minutes. EEG data were recorded via a BioSemi ActiveTwo EEG system at a sampling rate of 16,384 Hz.

Click trains of 100 µs with alternating polarity were presented at an average rate of 40 Hz. The timing of the clicks was randomized according to a pseudorandom Poisson process, as done before (Maddox & Lee, 2018; Polonenko & Maddox, 2024). Triggers marked the beginning and end of each 20 second trial and were created through the soundcard and converted to TTL through a custom trigger box (Maddox, 2020). The stimuli were presented diotically at 65 dB peSPL, which is quieter than typically done in clinical paradigms but was done to reflect more typical everyday listening levels. This level also stimulates the cochlea in a different part of the dynamic range than the higher intensity presented in Experiment 2 to investigate whether the different levels may reveal different relationships with the speech perception outcomes. Two runs of 4,800 sweeps (totaling 9,600 sweeps) were collected for each participant and took approximately four minutes to complete. Responses from the high forehead (electrode Fz) were analyzed and referenced to the average of electrodes placed on both earlobes.

#### 2.1.3 ABR Analysis and Peak-Picking

Custom Python scripts that used the MNE module (Gramfort et al., 2013) were used to preprocess and analyze the brainstem responses. Raw EEG was filtered between 150 and 2000 Hz using a first-order Butterworth filter and 5 Hz wide second-order infinite impulse response notch filters at odd multiples of 60 Hz. Epochs were created for each 20 second trial that included 1 second before and after the stimulus presentation for a total of 22 seconds. Brainstem responses were derived by cross-correlating the EEG epochs with similarly created zero-padded impulse sequences of clicks (a pulse train regressor created by placing an impulse at the time of each click and concatenating 1 second of zeros before and after the pulses), which was done in the frequency domain for efficiency. Each trial was weighted by the inverse of its variance divided by the sum of variances of all trials to create a Bayesian-like weighted average response (Polonenko & Maddox 2019, 2021, 2022, 2024a) so that noisy trials contributed less to the average.

ABR waveforms were analyzed by measuring peak amplitudes and latencies. Two separate examiners (M.J.P. and J.C.) labeled the peaks for waves I, III, and V for all participants so that consistency of wave labeling could be assessed. An intraclass correlation analysis (ICC) was performed using the psych package (Revelle, 2024) in R statistical software (R Core Team, 2022) using a two-way mixed effects model with “single fixed rater” units. The ICC analysis revealed excellent (ICC3 > 0.90) absolute agreement between the two raters for all metrics except a moderate agreement for wave I latency (ICC3 = 0.52), indicating reliable metrics of peak amplitudes and latencies.

#### 2.1.4 Competing Talker Task

Sentences for the speech-in-noise perception test used in this study were generated in Matlab (The MathWorks, Inc.) using the Boston University Gerald (BUG) corpus, which consists of 40 independent words that can be formed into syntactically correct but unpredictable sentences (Kidd et al., 2008). “Target” sentences were created using words from a male speaker with an F0 of ∼100 Hz. “Distractor” sentences were initially created using words from the same male speaker to control for talker idiosyncrasies affecting identification performance. Then the F0 of the Distractor sentences was shifted upward six semitones to ∼141 Hz using the “Change semitones” function within the Praat Vocal Toolkit (Boersma & Weenink, 2022). This manipulation provided participants with subtle F0 differences to segregate Target from Distractor. Although the use of a competing talker masker introduces informational masking effects and engages central auditory resources, this type of masker is much more representative of everyday listening than are purely energetic noise maskers.

The talker with an F0 of ∼100 Hz was the target talker on all trials, as pilot testing revealed that most listeners performed near chance levels when the target talker was varied throughout trials. One Target and one Distractor sentence were combined to create an audio file for each trial. The signal-to-noise ratio between target and masker was 0 dB such that intensity cues could not be used to differentiate the sentences. Audio file levels were scaled to 70 dB SPL.

The task was created in Gorilla Experiment Builder (www.gorilla.sc; Anwyl-Irvine et al., 2020). Participants were seated in a sound-attenuating booth and heard Target and Distractor sentences on each trial played diotically. The word “AND” spoken by the target talker was played at the beginning of each trial to cue listeners as to which talker (e.g., which F0) they should attend to during the trial. The Distractor sentence began 200 milliseconds prior to the Target sentence such that the words in each sentence overlapped somewhat but not completely, allowing each sentence to be perceived as an auditory stream rather than isolated, overlapping words. The Distractor, rather than the Target, sentence was chosen to begin first so that participants could not lock onto the Target immediately after the “AND” cue was played; their attention was first disrupted by the onset of the Distractor. Both sentences were presented in the same spatial location (0° azimuth) and therefore F0 was the primary cue available for participants to identify the Target sentence.

Participants heard sentences from two conditions: 1) control, in which the sentences contained their normal frequency information, and 2) high-pass filtered, in which F0 and the first eight harmonics were removed (0-800 Hz). As the target talker had an F0 around 100 Hz, high-pass filtering at 800 Hz removed the first 8 harmonics, leaving primarily unresolved harmonics that do not produce clear cochlear excitation peaks. Participants thus need to place greater reliance on pitch specifically coded by auditory nerve phase-locking to differentiate between the talkers and identify the target sentence (Oxenham, 2012). This task thus allows for a more thorough understanding of the role of subcortical neural function in speech-in-noise perception among normal hearing young adults than similar tasks used in previous research. High-pass filtered stimuli were created in Praat (Boersma & Weenink, 2022) by using a stopband from 0-800 Hz with 100 Hz smoothing.

To complete the competing talker task, participants first set the computer volume to a comfortable level while listening to control sentences. They then performed practice trials to familiarize themselves with the procedure, first with three trials in which control Target sentences were presented alone and then five trials in which control Target and Distractor sentences were presented simultaneously. Participants clicked on the words they heard in the Target sentence after each trial and feedback was provided. Performance on practice trials was not included in the data analysis.

Following practice, participants completed 30 test trials containing distinct sentences per condition divided into two blocks of 15 sentences each. The blocks containing control sentences were presented first followed by the blocks of high-pass filtered sentences. Sentence order was randomized within each block for each participant. The task took participants about 25 minutes on average to complete, including setting the computer volume and performing practice trials. Sentences were scored as the percentage of correctly identified words per condition. The 150 individual words were used as the scoring metric instead of the 30 whole sentences to increase the number of data points for each participant.

*2.1.5 Cognitive Task*

Participants performed a forward and backward digit span task to assess their short-term working memory. Digit span was chosen over another widely used measure, reading span, to limit any potential effects of a listener’s vocabulary on the task. The task was created in Gorilla Experiment Builder (www.gorilla.sc; Anwyl-Irvine et al., 2020) and consisted of sequences of digits presented visually so that individual differences in auditory perception did not influence the cognitive task (Smith & Pichora-Fuller, 2015). One digit was presented at a time and participants were asked to remember the digits and type them in a response box at the end of the digit sequence. In the “forward” condition, participants reported the digits in the order they were presented. In the “backward” condition, participants reported the digits in the reverse order of how they were presented.

Each condition began with three practice trials in which participants saw sequences of three digits and reported them according to the rules of the condition. Performance on the practice trials was not scored. Following practice, the test trials began with sequences of three digits and increased in sequence length to four, five, six, and seven digits. Three trials were presented per digit sequence length per condition. All participants completed the forward condition prior to completing the backward condition. Each trial was scored as “correct” if participants correctly reported all digits in the sequence in order (forward or backward), and “incorrect” if not. The percentage correct on the forward and backward conditions were averaged together to create one composite digit span score for each participant. The digit span task took approximately six minutes to complete.

#### 2.1.6 Experiment 1 Procedure

Participants were seated in a single-walled electrically shielded booth and first underwent the hearing screening. They then used a computer to perform the competing talker task, during which stimuli were presented through Sennheiser HD 650 open-back headphones, followed by the visual digit span task. The participant was then moved to a reclining chair in the booth to minimize muscle artifact for ABR recording. Clicks to elicit ABRs were presented through ER-2 insert earphones (Etymotic, Elk Grove Village, IL) coupled to an RME digiface USB soundcard at a sampling rate of 48 kHz. Participants were instructed to remain still and relax while they heard the clicks. Other EEG data were collected following ABRs and are not presented here. The total time for the measures described in Experiment 1 was one hour on average. Participants were allowed to take breaks both within and between measures and thus the total experiment time varied for each individual.

#### 2.1.7 Data Analysis

All statistical analyses were performed in R software (R Core Team, 2022). Although the competing talker task was designed to be sensitive to AN function, wave V metrics were included in statistical models in addition to wave I metrics to account for any effects of upstream subcortical processing deficits on participants’ speech-in-noise perception scores. Differential ABR metrics were also calculated and included in statistical analyses to normalize variability in wave amplitudes and latencies due to factors unrelated to neural physiology, such as individual differences in cochlear length and head size (Dehan & Jerger, 1990; Don et al., 1993). These included the ABR wave I:V ratio, calculated by dividing the amplitude of wave I by the amplitude of wave V, and the latency differences between wave V and wave I, calculated by subtracting the latency of wave I from that of wave V, for each participant.

A series of linear regressions were conducted to test independent hypotheses. First, we examined the effects of ABR wave I and V amplitudes and latencies on competing talker task scores in the high-pass and control conditions. Next, we examined the effects of normalized ABR wave metrics (ABR wave I:V ratio and wave I-V latency difference) on competing talker task scores from each of the two conditions. We then examined the potential influence of cognitive ability, assessed by the digit span task, on competing talker task performance.

### 2.2 Results

#### 2.2.1 Competing Talker Task Performance Is Variable Across NH Participants but Not Conditions

In the competing talker task, participants focused on a target talker while ignoring a competing talker. Figure 1 shows the performance of each participant for the two conditions, as well as the boxplot distributions (Shapiro-Wilk test *p* < 0.01, indicating the scores are not normally distributed). Chance performance was 12.5% correct and all participants performed above chance. While there was only a small change in performance across conditions (median difference = −2%; paired Wilcoxon signed-rank test, *V*(28) = 278, *p* = 0.194), there was a wide range in scores across participants. Performance in the control condition, which contained no high-pass filtering manipulation, ranged from 30.0% to 99.3% correct (median = 89.3% correct, interquartile range = 8.0%). Performance in the high-pass filtered condition, in which both talkers’ voices were manipulated to increase reliance on AN pitch encoding to segregate the competing talkers, ranged from 34.7% to 98.0% correct (median = 84.7% correct, interquartile range = 12.7%). Thus, despite normal audiometric thresholds, participants varied in their ability to correctly identify words spoken by a target talker in the presence of a competing talker. Next, we investigated whether ABR metrics and cognitive factors could explain some of this inter-participant variation in performance.

**Figure 1.**
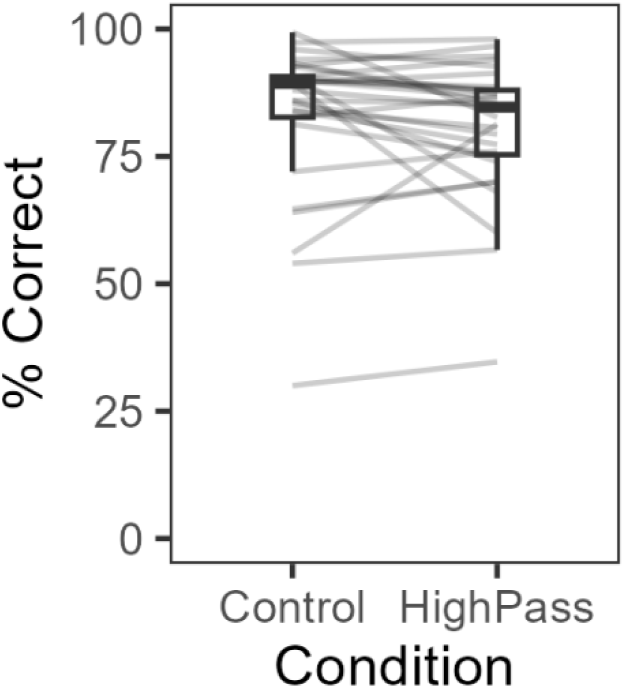
Experiment 1 percent correct word recognition scores on the competing talker task. The high-pass condition contained stimuli that had been high-pass filtered to minimize the contribution of cochlear pitch encoding, such that participants needed to primarily rely on pitch coded by auditory nerve phase-locking to differentiate between the target and distractor talkers. Stimuli in the control condition were not filtered. The top and bottom of each box are the 75th and 25th quartiles, respectively, and the line in the middle of each box represents the median. Whiskers indicate 1.5 times the interquartile range. Horizontal lines between the box plots connect an individual participant’s performance on each condition.

#### 2.2.2 Click ABR Metrics Do Not Predict Performance on The Competing Talker Task

Figure 2a shows individual participant and grand average ABR waveforms from this experiment. The regression analyses revealed that neither ABR wave amplitudes or latencies, nor normalized ABR metrics of I/V amplitude ratio and I-V inter-wave latency, significantly predicted performance on either condition of the competing talker task (p > 0.05 for all relationships; see Table 1). Surprisingly, even ABR wave I amplitude or latency did not significantly relate to participants’ scores on the high-pass filtered condition, which was designed to be particularly sensitive to AN function. Figure 2 also shows scatterplots of the relationships between ABR wave I and V amplitudes (panel B) and latencies (panel C) and performance on the high-pass filtered condition of the competing talker task.

**Figure 2.**
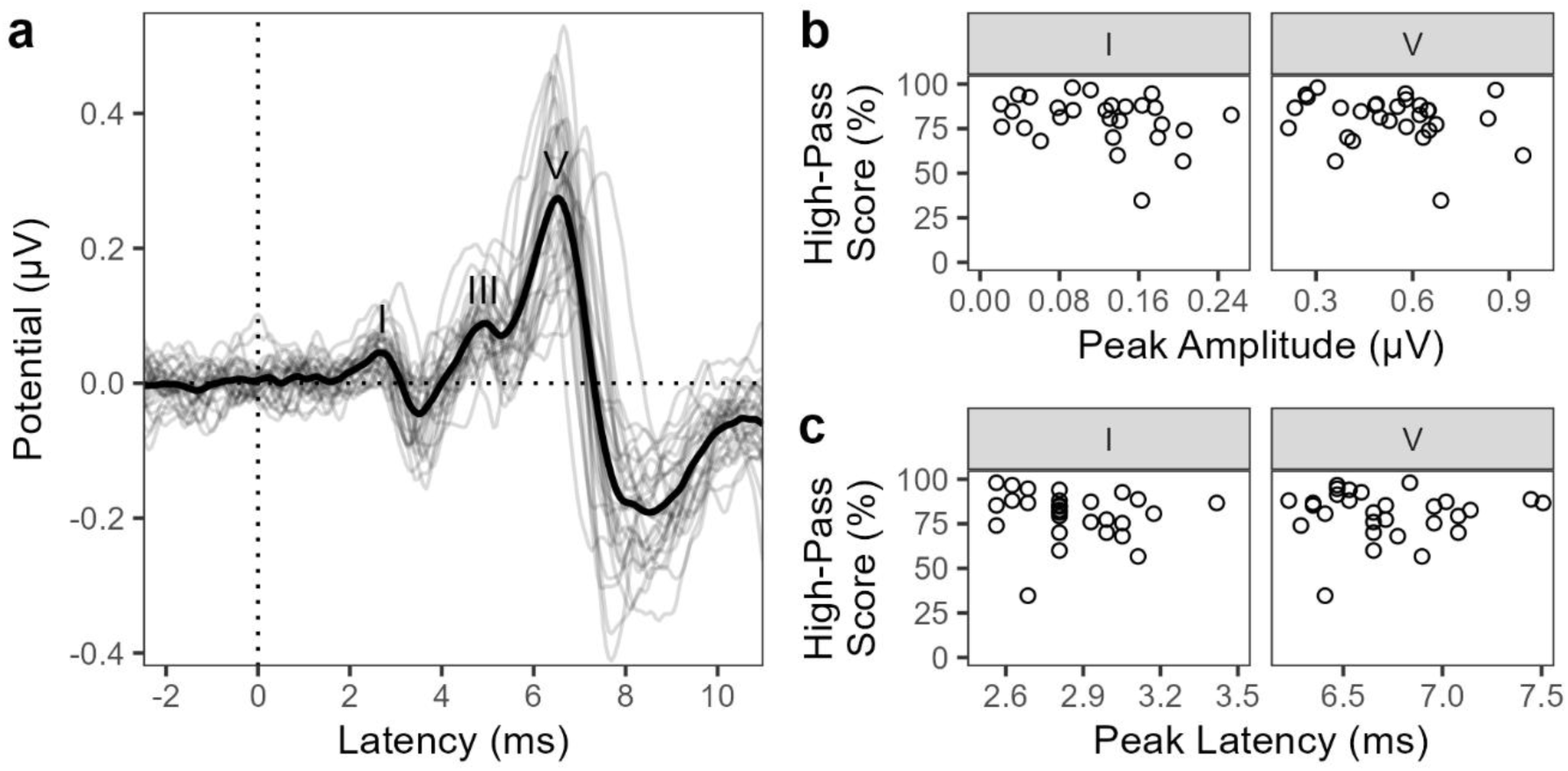
(A) Experiment 1 ABR waveforms. Waves I, III, and V are labeled. The thick, black line denotes the grand average across participants. Each gray line represents an individual participant’s waveform. Scatterplots show Experiment 1 relationships between word recognition scores on the high-pass filtered condition of the competing talker task and ABR wave I and wave V (B) amplitude and (C) latency. Each circle represents data from one participant.

**Table 1.** Results of linear regression models from Experiment 1 describing performance on the control and high-pass filtered conditions of the competing talker task as a function of ABR metrics and cognitive digit span task scores.

| **Dependent Variable** | **Model #** | **Independent Variable** | **Estimate ($\beta$)** | **Standard Error** | ***t*** | ***p*** |
| --- | --- | --- | --- | --- | --- | --- |
| **Control Condition Word Identification Scores** | 1 | Wave I Amplitude | -39.63 | 55.25 | -0.72 | 0.48 |
|  | 1 | Wave I Latency | -12.99 | 19.21 | -0.68 | 0.51 |
|  | 1 | Wave V Amplitude | -3.29 | 20.48 | -0.16 | 0.87 |
|  | 1 | Wave V Latency | 2.73 | 11.68 | 0.23 | 0.82 |
|  | 2 | Wave I:V Ratio | -21.66 | 24.93 | -0.87 | 0.39 |
|  | 2 | Wave I-V Latency Difference | 4.58 | 10.81 | 0.42 | 0.68 |
|  | 3 | Digit Span Score | 0.12 | 0.16 | 0.73 | 0.47 |
| **High-Pass Filtered Word Identification Scores** | 4 | Wave I Amplitude | -61.31 | 44.67 | -1.37 | 0.18 |
|  | 4 | Wave I Latency | -21.97 | 15.53 | -1.41 | 0.17 |
|  | 4 | Wave V Amplitude | -18.36 | 16.56 | -1.11 | 0.28 |
|  | 4 | Wave V Latency | 0.29 | 9.44 | 0.03 | 0.98 |
|  | 5 | Wave I:V Ratio | -22.51 | 21.68 | -1.04 | 0.31 |
|  | 5 | Wave I-V Latency Difference | 4.94 | 9.40 | 0.53 | 0.60 |
|  | 6 | Digit Span Score | 0.11 | 0.14 | 0.75 | 0.46 |

#### 2.2.3 Digit Span Scores Also Do Not Predict Performance on the Competing Talker Task

Composite digit span scores ranged from 23.3% to 93.3% correct, with an average score of 57.7% correct, demonstrating substantial variance in working memory scores among our young adult participants. Still, the linear regression analyses revealed no significant effects of working memory ability on competing talker task scores in the control (*p* = 0.47) or the high-pass filtered condition (*p* = 0.46). Figure 3 shows the relationship between composite digit span scores and performance on the high-pass condition of the competing talker task, the condition of most interest in this experiment.

**Figure 3.**
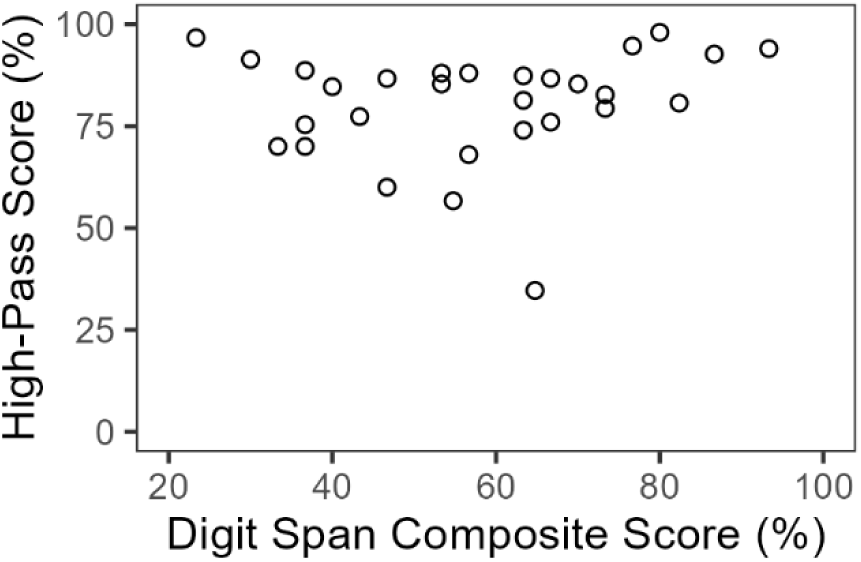
Experiment 1 relationship between participants’ composite scores on the digit span task and performance on the high-pass filtered condition of the competing talker task. Each circle represents data from one participant.

In this experiment, clicks were presented at a lower level than in clinical exams to emulate more typical listening levels. Yet, ABR metrics did not predict scores on a competing talker task that was presented at a similar level. In addition, working memory ability (as assessed by digit span scores) did not significantly relate to performance on the competing talker task.

Table 1 shows the results of these statistical models. A second experiment was therefore conducted using clinical parameters for ABR collection and tests of cognitive function in additional domains that are important for perceiving speech in challenging listening scenarios, such as attention, cognitive flexibility, and inhibitory control.

## 3. Experiment 2

### 3.1 Materials and Methods

#### 3.1.1 Participants

Thirty-one young adults aged 18-27 years (average age 21.69 years), distinct from those who participated in Experiment 1, completed Experiment 2. Some participants were recruited through advertisements in the Buffalo area and through the University at Buffalo Participate in Research Portal and completed the study for pay. Others were recruited through a department participant pool and completed the study for course credit. All participants were verified to have normal hearing thresholds (≤ 20 dB HL) from 250-8000 Hz. No participants exhibited hearing threshold asymmetries across ears, defined as threshold differences greater than 10 dB for two or more tested frequencies.

#### 3.1.2 Auditory Brainstem Response Recording

ABRs were recorded using Intelligent Hearing Systems Smart EP. In contrast to the parameters used in Experiment 1, Experiment 2 used the stimuli and the electrode configuration typically used for clinical assessment. The non-inverting electrode was placed at the top of the forehead and the ground electrode just above the center of the eyebrows. Inverting electrodes were placed on the mastoids. Electrode impedance levels were consistently kept below 3.0 kΩ, and electrodes were adjusted and/or replaced if necessary. Electrode placement took approximately five minutes. ABRs were recorded while participants passively listened to clicks of 100 µs duration with alternating polarity presented at 90 dB nHL. A presentation rate of 11.1 clicks per second was used to maximize the wave I response. Two runs containing 2048 sweeps, totaling 4096 sweeps, were completed for each ear. Collection of ABR responses took approximately 10 minutes.

#### 3.1.3 ABR Analysis and Peak-Picking

Two separate examiners (J.C. and I.K.) labelled the peaks for waves I, III, and V for data from all participants. Smart EP calculated the wave amplitudes and latencies for each run based on this peak-picking. Consistency in wave labelling between examiners was assessed via an ICC analysis, as described in Experiment 1. There was excellent absolute agreement between the two raters (ICC3 > 0.90) on all metrics.

The wave amplitudes and latencies of the two runs from each ear were averaged together. Values from each ear were then averaged to create a bilateral measure of wave I and Wave V amplitudes and latencies, as the competing talker task was presented binaurally. Normalized ABR metrics of the ABR wave I/V ratio and the latency difference between waves I and V were again calculated for participants in this experiment.

#### 3.1.4 Competing Talker Task

Participants first set the volume to a comfortable listening level and completed the same eight practice trials of control sentences and 30 test trials of high-pass sentences as in Experiment 1. High-pass filtered sentences were divided into two blocks and presented in random order. Participants did not perform test trials of the control sentences to limit potential effects of task adaptation prior to high-pass filtered sentence presentation. All other task parameters in Experiment 2 were identical to those used in Experiment 1. The task, including volume-setting and practice, took participants about 20 minutes on average to complete.

#### 3.1.5 Cognitive Tasks

Participants used an iPad to complete three assessments in the NIH Cognitive Toolbox V3 (Toolbox Assessments, Inc., 2023) each evaluating subsets of cognitive function that are important for speech-in-noise perception. All tasks provided participants with simultaneous verbal and written instructions. Practice trials were administered to ensure that participants understood each task. Performance on practice trials was not included in the scores for any test.

The Dimensional Change Card Sort (DCCS) test assessed attention and cognitive flexibility, the ability to switch cognitive or perceptual strategies to adapt to varying circumstances. The participant was shown two cartoon pictures at the bottom of the screen that were different shapes and colors (for example, a white rabbit and a brown boat). A third picture, the “target,” (for example, a brown rabbit) then appeared in the middle of the screen. The participant was instructed to select the picture at the bottom of the screen that matched either the color or shape of the target picture, cued by the word “color” or “shape” appearing on the screen with simultaneous audio of that word. If instructed to match based on “color,” they would click on the picture on the bottom of the screen that matched the color of the target picture (the brown boat). If instructed to match on “shape,” they would click on the picture on the bottom of the screen that matched the shape of the target picture (the white rabbit). Participants were told to respond as quickly as possible without making mistakes. Two practice rounds contained trials with instructions to match on color and one practice round contained trials instructing to match on shape. During the test round, trials varied between matching on shape or color. Scores were automatically calculated and were comprised of metrics related to both task accuracy and reaction time. The task took approximately four minutes to administer.

The Flanker Inhibitory Control and Attention (“Flanker”) test assessed the ability to selectively attend to a target stimulus and inhibit attention to other stimuli. A row of five cartoon fish was shown on each trial. Each fish had an arrow overlaid on them, pointing to either the left or the right. Participants were instructed to focus on the direction that the arrow on the middle fish was pointing and to ignore the other fish. They responded by clicking the button on the bottom of the screen that corresponded to the direction (left or right) that the arrow on the middle fish was pointing and were told to respond as quickly as possible. The task contained forty trials. On “congruent” trials, the arrows on all fish pointed in the same direction. On “incongruent” trials, the arrow on the middle fish pointed the opposite direction of the arrows on the surrounding fish. Scores were automatically calculated and were comprised of metrics related to both task accuracy and reaction time. The task took approximately three minutes to administer.

The List Sorting Working Memory (LSWM) test assessed working memory. Sequences of animal or food items were presented on each trial. A picture of the item and the written name of the item (e.g., “dog,”) were displayed on the screen at the same time as the name of the item was presented via audio. One item was presented at a time. The task began with the “one list” section, during which either only animal or food items were presented in each sequence. A chime at the end of each sequence prompted the participant to verbally state the presented items in size order from smallest to largest. Participants then completed the “two list” section, in which each sequence contained both food and animal items. At the end of each trial in this section, the participant verbally stated the presented food items from smallest to largest, followed by the presented animal items from smallest to largest. The first test trial in each section contained a four-item sequence. The sequences increased in length with each trial until the sequence contained seven items. The experimenter manually scored the task, selecting a 1 on a keyboard if the participant correctly sorted the entire sequence and 0 if they did not. Age-adjusted scores were automatically calculated. The task took approximately seven minutes to administer.

#### 3.1.6 Experiment 2 Procedure

Participants completed tasks for Experiment 2 in a double-walled sound attenuating booth. They first underwent the hearing screening followed by ABR collection. ABR stimuli were presented through ER-2 insert earphones (Interacoustics, Eden Prairie, MN) and the chair was reclined during recording to minimize muscle artifact. Participants were told to sit still and relax while they heard the clicks. They then used a computer in the booth and Sennheiser HD 650 open-back headphones to complete the competing talker task. Lastly, participants exited the booth and sat at a table in a quiet room to complete the NIH Cognitive Toolbox tasks. The total time for the measures described in Experiment 2 was about one hour and ten minutes on average. As in Experiment 1, participants were allowed to take breaks and thus the total experiment time varied for each individual.

#### 3.1.7 Data Analysis

Statistical analyses were performed in R software (R Core Team, 2022). Linear regression models were conducted to investigate the effects of 1) ABR wave I and V amplitudes and latencies and 2) normalized ABR metrics on participants’ performance on the competing talker task. A third linear regression model examined whether scores on the three cognitive tests significantly predicted competing talker task performance.

### 3.2 Results

#### 3.2.1 Competing Talker Task

Scores on the high-pass filtered condition, the only condition tested in this experiment, ranged from 30.0% to 93.3% correct (mean = 69.5% correct). The distribution of scores is shown in panels B and C of Figure 4. Average scores were lower in Experiment 2 than in Experiment 1 (see Figure 1), possibly because participants in Experiment 1 had greater task familiarization due to hearing blocks of control sentences prior to high-pass filtered sentences. However, participant populations also differed between the experiments and it is likely that participant variables and other experimental factors contributed to score differences between experiments.

**Figure 4.**
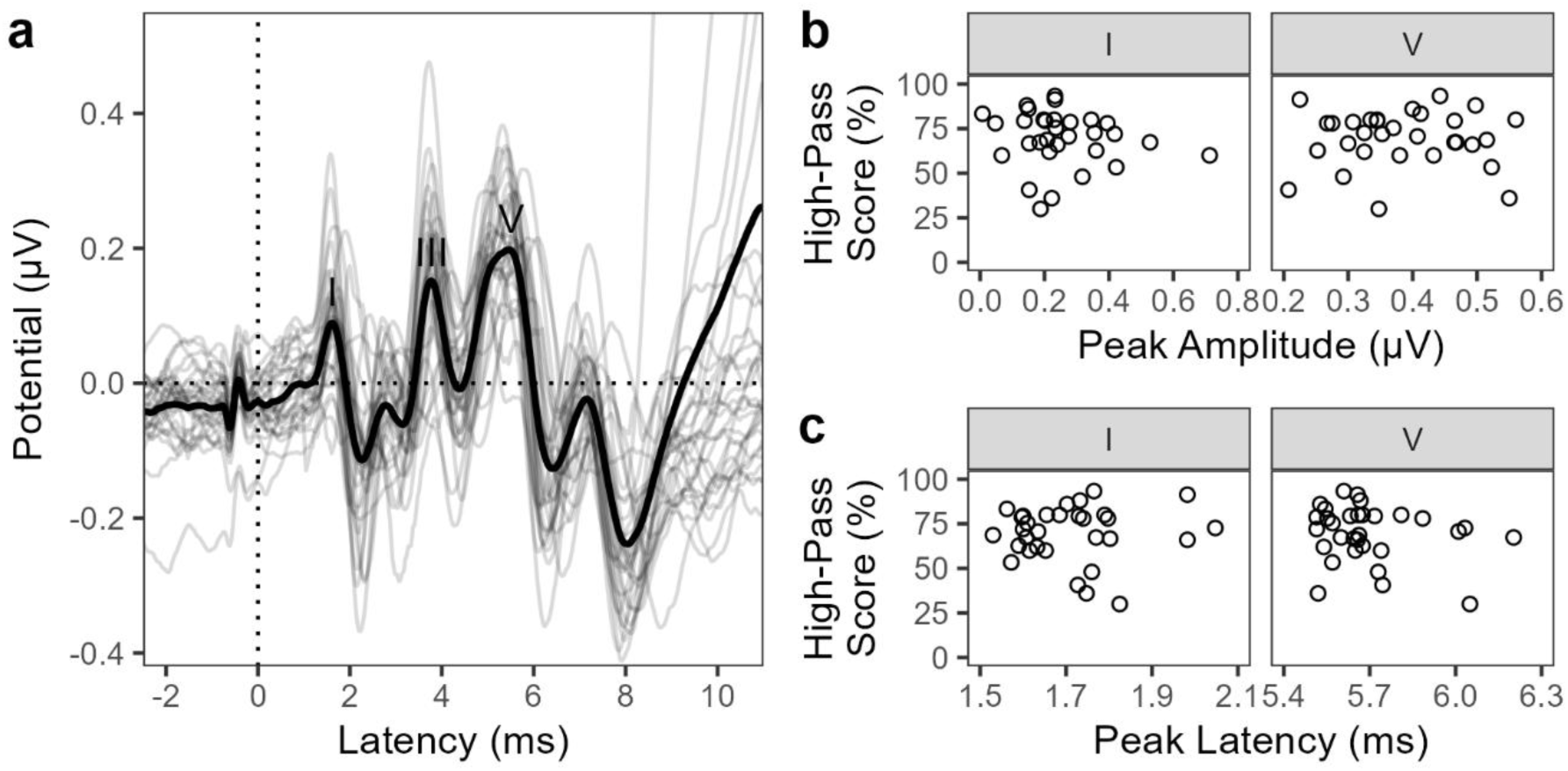
(A) Experiment 2 ABR waveforms. Waves I, III, and V are labeled. The thick, black line denotes the grand average across participants. Each gray line represents an individual participant’s waveform. Scatterplots show Experiment 2 relationships between word recognition scores on the high-pass filtered condition of the competing talker task and ABR wave I and wave V (B) amplitude and (C) latency. Each circle represents data from one participant.

#### 3.2.2 Clinical ABR Metrics Do Not Predict Performance on the Competing Talker Test

Figure 4a shows individual participant and grand average ABR waveforms from this experiment. Although we again observed variability in ABR metrics among participants, no significant relationships were found between ABR wave I and V amplitudes or latencies, nor normalized ABR metrics, and performance on the high-pass filtered condition of the competing talker task (p > 0.05 for all relationships; see Table 2). Figures 4a and 4b illustrate the relationships between ABR wave I amplitudes and latencies and high-pass filtered competing talker task scores from this experiment.

**Table 2.** Results of linear regression models from Experiment 2 describing performance on the competing talker task as a function of ABR metrics and cognitive task scores. DCCS = Dimensional Change Card Sort test. Flanker = Flanker Inhibitory Control and Attention test. LSWM = List Sorting Working Memory test. * = *p* is significant at a level of 0.05.

| Dependent Variable | Model # | Independent Variable | Estimate ($\beta$) | Standard Error | $t$ | $p$ |
| --- | --- | --- | --- | --- | --- | --- |
| **High-Pass Filtered Word Identification Scores** | 1 | Wave I Amplitude | -27.72 | 21.02 | -1.32 | 0.20 |
|  | 1 | Wave I Latency | -20.02 | 32.85 | -0.61 | 0.55 |
|  | 1 | Wave V Amplitude | -15.94 | 31.30 | -0.51 | 0.62 |
|  | 1 | Wave V Latency | -32.19 | 24.90 | -1.29 | 0.21 |
|  | 2 | Wave I:V Ratio | -4.98 | 7.24 | -0.69 | 0.50 |
|  | 2 | Wave I-V Latency Difference | -18.24 | 23.57 | -0.77 | 0.45 |
|  | 3 | DCCS Score | -0.29 | 0.25 | -1.17 | 0.25 |
|  | 3 | Flanker Score | 0.68 | 0.26 | 2.65 | 0.01* |
|  | 3 | LSWM Score | -0.14 | 0.23 | -0.60 | 0.56 |

#### 3.2.3 Competing Talker Performance is Predicted by Cognitive Inhibitory Control but Not Cognitive Flexibility or Working Memory

The regression analysis to examine the influence of cognitive task scores on speech-in-noise perception performance revealed that Flanker Inhibitory Control and Attention (Flanker) test scores significantly predicted participants’ ability to identify target sentences in the high-pass condition of the competing talker task (*p* = 0.01). Scores on the Dimensional Change Card Sort (DCCS) test (*p* = 0.25) and the List Sorting Working Memory (LSWM) test (*p* = 0.56) were not significant predictors. Multicollinearity was observed in this regression model, as scores on the Flanker and DCCS were correlated. Revising the models by first removing the Flanker test scores, then adding Flanker test scores back in and removing DCCS test scores, did not change the overall results (Flanker: *p* = 0.02; DCCS: *p* = 0.76 from revised models). The relationships between cognitive task scores and high-pass filtered competing talker task scores are shown in Figure 5. The results of the regression models run in Experiment 2 are shown in Table 2.

**Figure 5.**
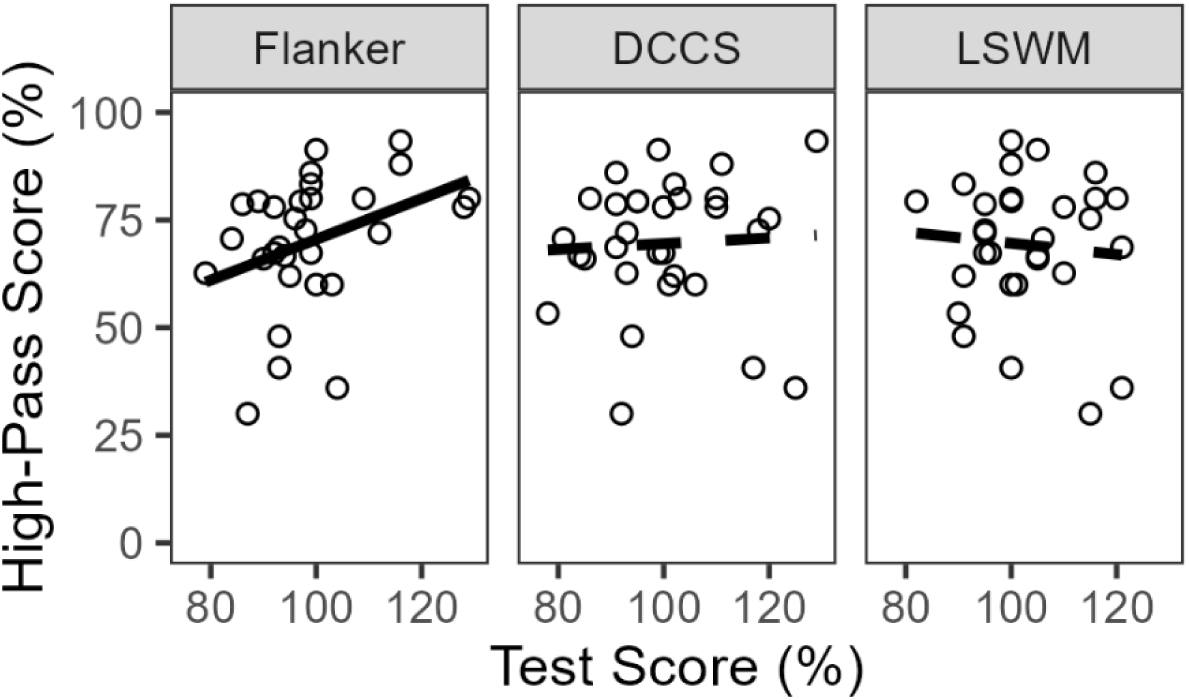
Relationships between participants’ scores on the Flanker Inhibitory Control and Attention (Flanker) test, Dimensional Change Card Sort (DCCS) test, and List Sorting Working Memory (LSWM) test and performance on the high-pass filtered condition of the competing talker task. Each circle represents data from one participant. The solid black line indicates the line of best fit for the significant relationship between Flanker test scores and competing talker task scores. The dashed black lines represent the lines of best fit for the non-significant relationships.

Follow-up correlation analyses were conducted to examine whether listeners with decreased ABR wave amplitudes or increased ABR wave latencies were those who performed better on the cognitive tasks, as a potential compensatory mechanism. No significant correlations were found between ABR metrics and scores on the Flanker, DCCS, and LSWM tests. It seems likely that the participants in this study did not have clinically significant deficits in AN and subcortical function that warranted the use of compensatory strategies.

## 4. Discussion

This study examined the extent to which auditory neural processing and cognitive function predicted young adults’ ability to attend to a target talker while ignoring a competing talker – a common listening situation that can be challenging for some individuals with normal hearing thresholds (Hind et al., 2011; Kamerer et al., 2022). We conducted two experiments using either moderate- or high-intensity click stimuli for ABR collection to evaluate whether different cochlear and neural activation levels correlated with scores on a competing talker task. Although ABR wave amplitudes and latencies varied among participants in both experiments, neither experiment found a relationship between ABR metrics and performance on the competing talker task. These findings, combined with the result that decreased neural processing did not correspond to a compensatory increase in cognitive task scores, suggest that young, normal hearing adults who do not necessarily report difficulty hearing in noise do not have deficits in AN and subcortical function significant enough to impact perceptual ability in noise.

The results of this research therefore provide a potential explanation for the mixed results of previous studies that have explored whether AN and subcortical function, as measured by ABR wave amplitudes and latencies, predicts speech-in-noise perception scores. For example, Grose et al., (2017), Prendergast et al., (2017a), and Smith et al., (2019) tested young adults with normal hearing thresholds and did not find relationships between click ABR metrics and scores on various tests of hearing in noise. However, other studies that *have* observed this relationship included participants from a wider age range (Mepani et al., 2020; Grant et al., 2020) and/or those with some degree of hearing loss (Bramhall et al. 2015; Valderrama et al., 2018). Therefore, individuals may need to be older, have greater risk factors for cochlear synaptopathy or other forms of neural degeneration, or clinically significant cochlear damage for ABR metrics to predict speech-in-noise perception performance.

This study also examined the influence of cognitive abilities on speech-in-noise perception scores of young, normal hearing adults. We assessed participants’ working memory in both experiments in this study, as holding information in memory is necessary to accurately follow everyday conversations. Participants performed a forward and backward digit span task in Experiment 1 and the NIH Cognitive Toolbox List Sorting Working Memory test in Experiment 2. However, scores on the working memory assessments did not predict performance on the competing talker task in either experiment. One potential explanation for this result is that the working memory load in the competing talker task was moderate. Although participants were required to hold a sentence in memory before responding, the sentences always consisted of five words in the predictable structure “[name] [verb] [number] [adjective] [noun].” This may have eased cognitive load through structural priming, by allowing participants to scaffold the words to this structure (Pickering & Ferreira, 2008).

Another potential explanation for the finding that working memory did not predict competing talker task scores is that working memory capacity only plays a substantial role if the task is sufficiently demanding. The six-semitone separation between talkers is a strong cue for speech stream segregation (Darwin et al 2003; Flaherty et al 2019) and may have reduced reliance on cognitive resources in our participants, who were individuals with optimal auditory perception. This could explain why most investigations that have found a positive relationship between working memory task performance and speech-in-noise perception scores included middle-aged and older participants and those with hearing loss, populations that would experience greater difficulty on a speech-in-noise perception task than the young, normal hearing adults used in this study. For example, Rudner et al. (2011) found that performance on a reading span task significantly predicted speech recognition in different types of noise among hearing aid users. Yeend et al. (2019) tested participants aged 30-60 years with hearing thresholds ranging from normal hearing to moderate loss and found that reading span task performance predicted a composite measure of speech-in-noise perception consisting of self-reported listening ability and scores on two speech identification tasks. Other studies that have tested both young and older adults found reading span performance to correlate with speech-in-noise identification scores among only the older adults (Füllgrabe & Rosen, 2016a; Vermeire et al., 2019). The current study adds to the literature suggesting that working memory ability does not play a significant role in the speech-in-noise perceptual abilities of individuals without sensory impairment or any known challenges hearing in noise. We also extend the working memory assessments supporting this conclusion to digit span and the List Sorting Working Memory test.

The second experiment in this study expanded the cognitive test battery beyond working memory to measures of cognitive flexibility, attention, and inhibitory control. Scores on the Flanker Inhibitory Control and Attention Test were found to significantly predict performance on the competing talker task. This is likely due to the similar perceptual demands of both tasks. In the Flanker test, participants were asked to focus on a visual target presented simultaneously with competing visual information. The competing talker task required the same perceptual processes, but in the auditory domain: attend to a target talker and ignore a distractor talker. The significant relationship between performance on the Flanker test and the competing talker task adds to the evidence that similar mechanisms underlie both visual and auditory attention (Shinn-Cunningham, 2008). This result also highlights the importance of examining predictors of auditory perception that tap into the same processes as the auditory task itself. Scores on cognitive tests whose demands were more distinct from those necessary for the speech-in-noise perception task used in this study did not significantly relate to task performance.

Likewise, the extent to which measures of auditory system function can explain individual differences in auditory perception may also depend on the similarity between the stimuli used to elicit auditory system responses and those used in the perceptual task. This investigation used click stimuli because their synchrony elicits robust wave I responses, the wave primarily of interest in this investigation. Although Experiment 1 recorded ABRs to clicks presented at similar levels as the stimuli in the competing talker task, cochlear and neural activation in response to clicks of any intensity differs from the activation elicited by speech. Speech is more temporally and spectrally dynamic than click trains even if the clicks are randomly presented. The recent development of a continuous speech ABR protocol (Polonenko & Maddox, 2021, 2024a) may provide more realistic measures of auditory processing that better predict speech perception. It has recently been established that continuous speech ABRs can be measured to more real-life paradigms that include multiple simultaneously presented talkers that are presented at conversational levels (Polonenko & Maddox, 2024b). Future experiments could determine if auditory processing metrics using multi-talker stimulus paradigms are more predictive of the variation in perception of the same stimuli, thus potentially making a more informative clinical measure of daily listening challenges.

The speech-in-noise perception task used in this study was created to be more sensitive to AN processing than are commonly used tests. Stimuli were high-pass filtered to remove the harmonics that create clear cochlear excitation peaks, requiring a listener to rely more on pitch coded by AN phase-locking to differentiate between two talkers with similar fundamental frequencies. However, cognitive, rather than neural, factors predicted task performance; considering that this test was designed to be specifically sensitive to deficits in AN function, this result further supports the conclusion that the young adults generally do not have clinically significant deficits in neural function, informing both past and future work using this participant population. Still, the significant relationship between performance on the high-pass condition of the competing talker task and Flanker test scores demonstrates the utility of this competing talker task for studies in young adults. This finding suggests that this task is tapping into real-world processing among listeners with normal hearing and demonstrates the utility of this task for future studies of listeners without audiometric loss.

This study has several limitations. First, the competing talker task contained only 30 trials. Identification of the words comprising the sentences, rather than identification of the full sentences themselves, were used as the outcome metric to increase the number of data points per participant. Still, a greater number of trials would have provided more data and allowed for analysis of sentence-level accuracy, which may have better related to working memory and other cognitive domains than did word-level accuracy. Second, each experiment used a different population of participants. This, in addition to participants hearing control and high-pass filtered sentences in Experiment 1 but only high-pass filtered sentences in Experiment 2, prevent comparison of how the ABRs elicited with different parameters related to performance on the competing talker task. We also chose to use digit span and List Sorting Working Memory rather than reading span or listening span tasks to minimize the influence of linguistic contributions on working memory scores. However, digit span and list sorting only tax working memory capacity, not the linguistic memory processing that is necessary for the competing talker task. A task such as reading span or listening span, which assess memory processing in addition to storage capacity (Daneman & Carpenter, 1980), may have been better suited to find a relationship between working memory performance and competing talker task scores.

Both experiments in this study used ABRs in order to isolate responses from the AN and subcortical nuclei. Yet, ABRs are neuron onset responses and not a measure of the phase-locking that is important for speech stream segregation by pitch cues. Frequency following responses (FFRs) might better assess neural contributions to performance on the competing talker task. However, FFRs do not have distinct neurogenerators and originate from both subcortical and cortical auditory neurons (Coffey et al., 2019). The competing talker task was designed to tax AN function, but AN phase-locking cannot be isolated with an FFR to directly compare to performance on this task. Speech-in-speech ABRs may allow for isolation of both AN responses and assessment of the dynamic neural function that is necessary for segregation of a target talker from a competing talker (e.g., Polonenko & Maddox, 2024b).

Lastly, this study did not examine the potential influence of extended high frequency (EHF) hearing (e.g., 9-16 kHz) on competing talker task scores. While the relevant energy in speech sounds is at lower frequencies, high frequency content can improve segregation of competing auditory sources (Monson et al., 2019) and improve speech-in-noise perception (Polspoel et al., 2022). Research has shown that young adults with normal hearing thresholds at standard frequencies (≤8 kHz) demonstrate substantial variability in EHF thresholds, and those with elevated EHFs are those who self-report challenges hearing in noise (Motlagh Zadeh, et al. 2019). The ABR metrics in the current investigation reflect cochlear function, but the click stimuli used to elicit ABRs did not range above 10 kHz due to ER-2 insert earphone frequency limitations. It is therefore possible that participants with poorer EHF processing were those who performed more poorly on the competing talker task. Collection of extended high frequency hearing thresholds in future work could provide additional information about the factors contributing to competing talker task performance.

## 5. Conclusion

Young adults with normal hearing thresholds demonstrate variability in ABRs to moderate and high intensity clicks, but individual differences in this population do not seem to be clinically significant enough to affect perception—even on a task specifically designed to be sensitive to AN function. Instead, a cognitive task with very similar demands to the competing talker task was the most significant predictor. These results may help provide insight into why many previous studies in young, normal hearing adults have not found a relationship between AN and subcortical function and speech perception in challenging listening environments. The participants in this study were not specifically recruited due to listening difficulties, noise exposure, or age, which may be factors necessary to find deficits in auditory processing that predict perceptual ability. Future work will replicate this research with a larger participant population who also vary more in their auditory system function, such as middle-aged and older adults. Assessing individuals with greater likelihood of exhibiting clinically relevant deficits in auditory perception will provide additional information about the role of decreased neural function, and potential compensatory cognitive mechanisms, in the ability to hear in background noise.

## Data Availability

The data that support the findings of this study are available at https://doi.org/10.5061/dryad.kprr4xhft.

## Acknowledgments

We thank PuiYii Goh and Isabel Herb for assistance with Experiment 1 data collection and Elizabeth Fish for assistance with Experiment 2 data collection. This work was funded by a College of Liberal Arts Social Sciences Seed Grant from the University of Minnesota (MJP) and an award from the Research and Creative Activities for Undergraduates Program at the University at Buffalo (MD).

